# *Drosophila melanogaster* females prioritise dietary sterols for producing high quality eggs

**DOI:** 10.1101/2021.06.04.447167

**Authors:** Brooke Zanco, Lisa Rapley, Joshua N Johnstone, Amy Dedman, Christen K Mirth, Carla M Sgrò, Matthew DW Piper

**Affiliations:** Monash University

## Abstract

Limiting calories or specific nutrients without malnutrition, otherwise known as dietary restriction (DR), has been shown to extend lifespan across a broad range of taxa. Our recent findings in *Drosophila melanogaster* show that supplementing flies on macronutrient-rich diets with additional cholesterol can extend lifespan to the same extent as DR. Macronutrient-rich diets drive high levels of egg production and in doing so deplete the mothers of somatic sterols that are essential for survival. Thus, DR may be beneficial for lifespan because it reduces egg production which in turn reduces the mother’s demand for sterols. If this is true, mothers must be prioritising their available sterols, whether from the diet or from their own bodies, to sustain high quality egg production. To test this, we measured the quality of eggs laid by mothers fed either cholesterol-sufficient or cholesterol-depleted diets. We found that even when the mother’s diet was completely devoid of cholesterol, high quality egg production persisted. Furthermore, we show that sterol-supplemented flies with long lives continue to lay high quality eggs that give rise to healthy offspring. Thus, in our assays, long life does not require a fecundity cost.

## Introduction

Changing the nutritional balance of an animal’s diet can extend lifespan in a manner that accounts for the full effects of dietary restriction (DR) (Mair *et al*., 2005; Lee *et al*., 2008; Simpson and Raubenheimer, 2007; Skorupa *et al*., 2008; Grandison *et al*., 2009; Solon-Biet *et al*., 2014, 2015; Regan *et al*., 2020). Specifically, across a broad range of species higher protein : carbohydrate (P:C) ratios are associated with short lifespan and high reproduction, while low P:C diets are associated with longer life, but lower levels of reproduction (Maklakov *et al*., 2008; Piper *et al*., 2011; Fanson and Taylor, 2012; Simpson *et al*., 2012, 2017). Our recent work provides a new nutritional explanation for this phenomenon by demonstrating that while P:C ratio does positively associate with female reproduction in *Drosophila*, it is the abundance of a different nutritional component, dietary sterols, that dictates length of life (Zanco *et al*., 2021). Importantly, the dietary P:C ratio manipulates the availability of sterols accessible for somatic maintenance, by either inhibiting or driving reproductive output, of which sterols are required for.

Based on our previous data, we proposed that in *Drosophila*, dietary P:C ratio is monitored to set the level of egg laying, while sterols, which determine length of life, are constitutively prioritised for reproduction (Zanco *et al*., 2021). Thus, when reproduction is low (on low P:C food) the diet supplies sufficient sterols to meet the relatively low demands for reproduction and this spares the mother from drawing on her own sterol reserves, which leaves her soma intact to sustain longer life. By contrast, on higher P:C diets that contain a proportionally low, inadequate, supply of sterols (e.g. high yeast food), the flies preferentially supply sterols to sustain reproduction at a faster rate than can be replenished from the diet. Thus, they draw on their own somatic reserves, which eventually depletes key tissues of sterols, reducing lifespan.

The constitutive prioritisation of limiting sterols to reproduction on both low and high protein food is consistent with the Nutrient Recycling Hypothesis, which argues that it is more advantageous for flies to maximise reproduction with whatever resources they have available, even if it means reducing maternal lifespan (Adler and Bonduriansky, 2014). This drive to meet the short term needs for reproduction is preferred because any strategy that involves protecting the soma to enhance future possible reproduction would rarely be beneficial in the wild due to the high risk of dying from extrinsic hazards (Adler and Bonduriansky, 2014). If true, we would expect that egg quality should not be compromised on sterol depleted diets.

An alternative hypothesis, the “Direct constraints” model of reproduction, can also explain our previous observations, but in this model, cholesterol is constitutively prioritised for use by the soma for maintenance rather than for reproduction (O’Brien *et al*., 2008; Tatar, 2017). This hypothesis argues that reproduction induces damage to the somatic tissues due to a multitude of factors. For instance, changes in metabolic activity during the reproductive phase can lead to an increase in oxidative damage while mating effort has been shown to induce immunosuppression (Nordling *et al*., 1998; Fedorka *et al*., 2004; Dowling and Simmons, 2009; Latta *et al*., 2019). Thus, restricted diets support longer lifespan because reproduction is low on these diets, meaning that even the limited dietary resources available are sufficient to repair damage. By contrast, on nutrient rich diets, elevated reproduction leads to an increase in damage above the levels that the diet can counter, and this shortens lifespan. If this were true in flies, when dietary yeast is increased, the resultant increase in P:C would cause greater reproduction-related damage than what the sterol supply could counter. In contrast, when dietary yeast is low, the fall in P:C ratio would lower reproduction to a level such that the damage it inflicts could be met by the small amounts of sterols that are provided in the food. Under this scenario, because limiting sterols are prioritised for somatic maintenance over reproduction, the higher P:C diets that elevate egg laying and shorten lifespan should be accompanied by the production of sterol depleted eggs, which compromises their viability (Heier *et al*., 2021).

Under both of these models, supplementing sterols in a high P:C (high yeast) diet should increase the mothers’ lifespan, which is what has been observed (Zanco *et al*., 2021), but they predict different effects on egg-to-adult viability. In the Nutrient Recycling Hypothesis (Adler and Bonduriansky, 2014), because the flies already prioritise sterols for reproduction, adding more to the diet would not markedly increase egg-to-adult viability, as it should already be high. Instead, the added sterol should create a surplus over what is required for egg production that mothers can use to preserve the soma to sustain longer life. In the direct constraints model (O’Brien *et al*., 2008; Tatar, 2017), sterol-supplemented flies would live longer because they would have increased their capacity to repair somatic damage, and because the mothers were already prioritising sterols for use by the soma, any excess from the addition of sterols to the diet should increase what is available for egg production and so we should see that egg-to-adult viability increases from low to high. Distinguishing between these possibilities is important as it gives us insights into the likely way in which nutrients and their depletion may cause death – the very key to understanding the beneficial effects of diet on lifespan. To distinguish between these possibilities, we manipulated the major nutritional determinant of lifespan (sterols) independently of the major nutritional determinant of egg production (P:C ratio) and monitored egg-to-adult viability and offspring development.

## Results

### Maternal sterol supplies are preferentially used to produce high quality eggs when dietary intake is limited

If egg quality is prioritised by mothers, then the egg-to-adult viability of eggs from young mothers should be sustained at a high level when fed either a cholesterol-sufficient or cholesterol-depleted diet. In contrast, if the flies prioritise somatic maintenance then we expect that egg-to-adult viability will be low for mothers on cholesterol-depleted diets, and high when they are fed a cholesterol-sufficient diet. To test this, we used our completely defined (holidic) diet (Piper et al., 2014) to manipulate cholesterol independently of all other nutrients. Mated females were placed on experimental diets containing either 0g/l or 0.3g/l (sufficient) cholesterol for 16 days. We scored the total number of eggs laid and egg-to-adult viability daily.

We found that while the number of eggs laid declined over time for flies on both sterol sufficient and depleted foods (Figure 1a), flies from both treatments were able to sustain maximum egg-to-adult viability for approximately 10 days (Figure 1b), at which point, egg laying had dropped to approximately four eggs per day for non-supplemented mothers (Figure 1a). This pattern of sustaining high egg-to-adult viability, which then drops off rapidly as egg production nears zero (as opposed to a slow linear decline in viability) is what we would predict if mothers prioritised egg quality over somatic maintenance.

**Figure 1.**
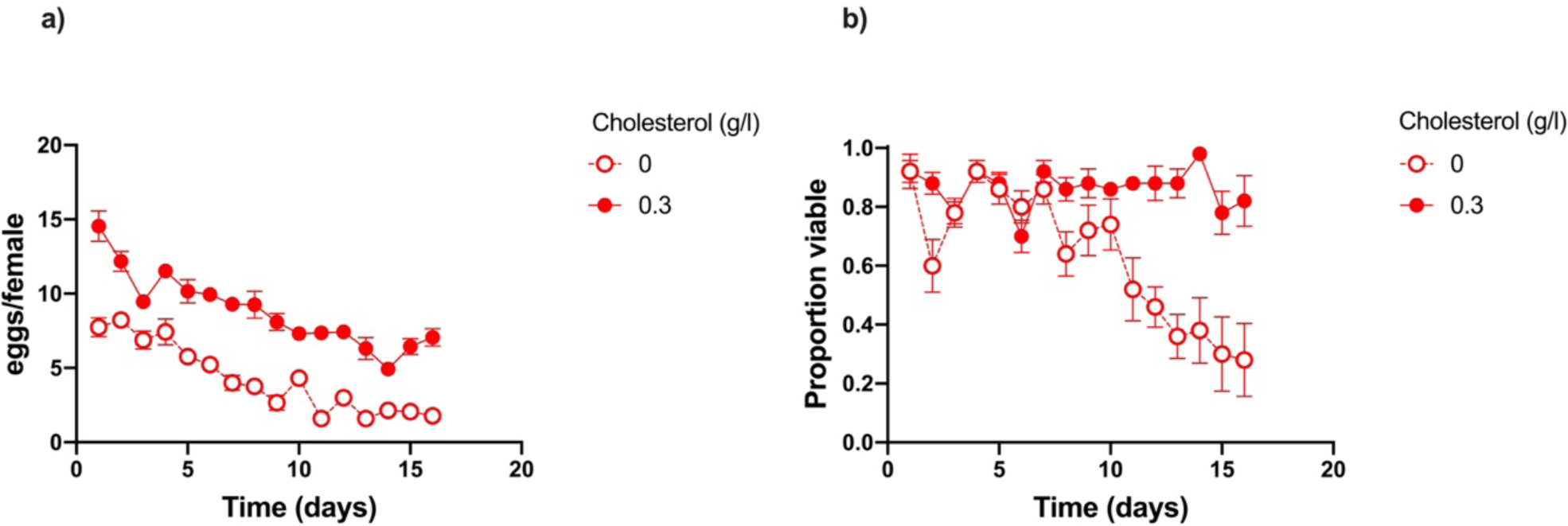
When flies were fed a fixed P:C ratio, removing cholesterol from the diet resulted in a sustained reduction in daily egg production (a). Egg-to-adult viability remained high across treatments until day 10, with no significant difference in mean viability when compared against the mean viability of all previous days (P > 0.05). After day 10 the total number of eggs laid by mothers fed 0g/l cholesterol dropped to 4 eggs per female and viability began to fall (a, b), at which point the mean viability on all successive days were significantly lower than the mean viability on day 10 (P < 0.05), except for day 11 which was not significantly different (P = 0.123). Each data point is the average (+/- s.e.) of eggs from five vials, each containing ten females (a). Viability (b) was assessed by transferring ten eggs into each of five replicate vials. Each data point is the average (+/- s.e.) of viable individuals across these vials.

While egg-to-adult viability is sustained, it is possible there is a cost during development for larvae from sterol depleted mothers. To assess this, we monitored egg-to-adult viability, egg-to-adult development time, and final adult body size of larvae that emerged from eggs laid on day 10 - the time point right on the verge of when mothers on the sterol-depleted food started to lay inviable eggs. As before, we found that there was no effect of maternal cholesterol on egg-to-adult viability, but interestingly, adding cholesterol to the larval diet did significantly increase the egg-to-adult viability of offspring, indicating a rescue of a sterol shortfall for development (Figure 2a). Larvae from eggs laid by sterol-supplemented mothers developed quicker than those from mothers whose diet did not contain sterols (Figure 2b), and they achieved a smaller final body size as adults than offspring from mothers without sterols (Figure 2c). Finally, supplementing the larval medium with cholesterol brought the development time and adult body size of offspring to the same level, irrespective of the mothers’ dietary sterol condition (Figure 2b, c). Interestingly, these values were intermediate to the differences found between treatments when larvae did not receive a cholesterol supplement. Together, these data show that mothers match egg production to the number of high-quality eggs they can produce, for ∼10 days after encountering a sterol-depleted diet. Furthermore, any effects of the mothers’ diet which modify egg-to-adult viability, offspring development time, and adult body size, are normalised to a common level when the larvae ingest sterol-replete food.

**Figure 2.**
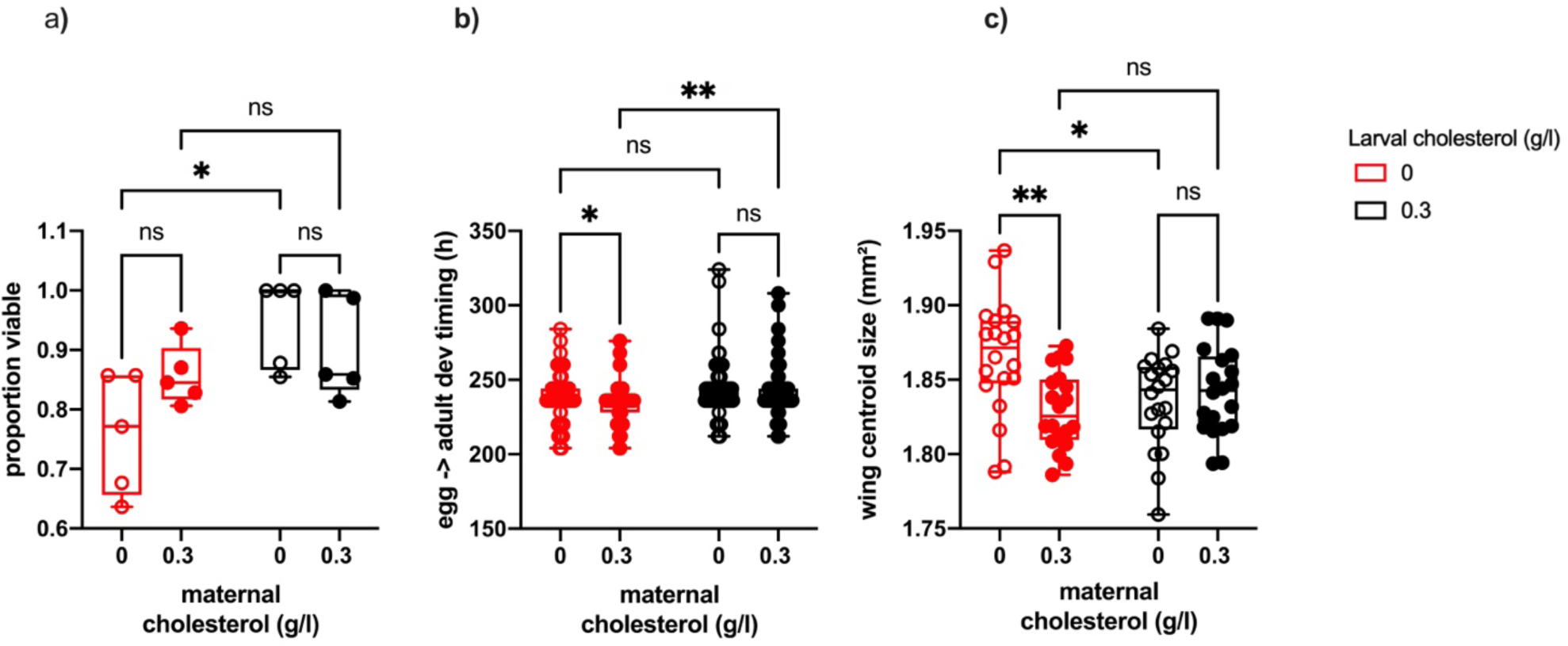
The amount of cholesterol present in either the maternal diet, larval diet or both, alters offspring fitness, however the direction of this variation differs between traits (a – c). Egg-to-adult viability was not significantly altered by maternal cholesterol levels, however, when mothers were fed a diet with 0g/l cholesterol, egg-to-adult viability increased significantly when the larval diet was supplemented with 0.3g/l cholesterol (a). Offspring developed significantly faster when mothers were fed 0.3g/l cholesterol (b), and their final body size (measured as wing centroid size) was significantly smaller (c). These effects disappeared when the larval diet was supplemented with 0.3g/l cholesterol (c, d). Ns = non-significant, * = < 0.05, ** = < 0.005, *** = < 0.001

### Maternal cholesterol levels do not significantly impact egg quality despite limiting maternal lifespan

The previous experiment used diets without sterols to assess the mothers’ commitment to producing high-quality eggs when faced with extreme nutrient stress. To test the extent of maternal sterol prioritisation under more natural conditions, we exposed flies to a standard sugar/yeast diet, which we have previously shown to be sterol limiting for female egg production and lifespan (Zanco *et al*. 2021), and supplemented it with sufficient sterols to overcome these limitations (0.3g/l cholesterol).

The same 10-day timepoint and parameters of egg quality were used as above. We found that there was no significant effect of either maternal or larval cholesterol supplementation on egg-to-adult viability, which remained at the highest level across all experimental conditions (Figure 3a). Furthermore, offspring development time (Figure 3b) was not affected by sterol addition (Figure 3c). In line with our earlier experiment, offspring achieved a smaller final body size when the maternal diet was supplemented with additional sterols, however here this only occurred when the larval diet was also supplemented with 0.3g/l of cholesterol (Figure 3c).

**Figure 3.**
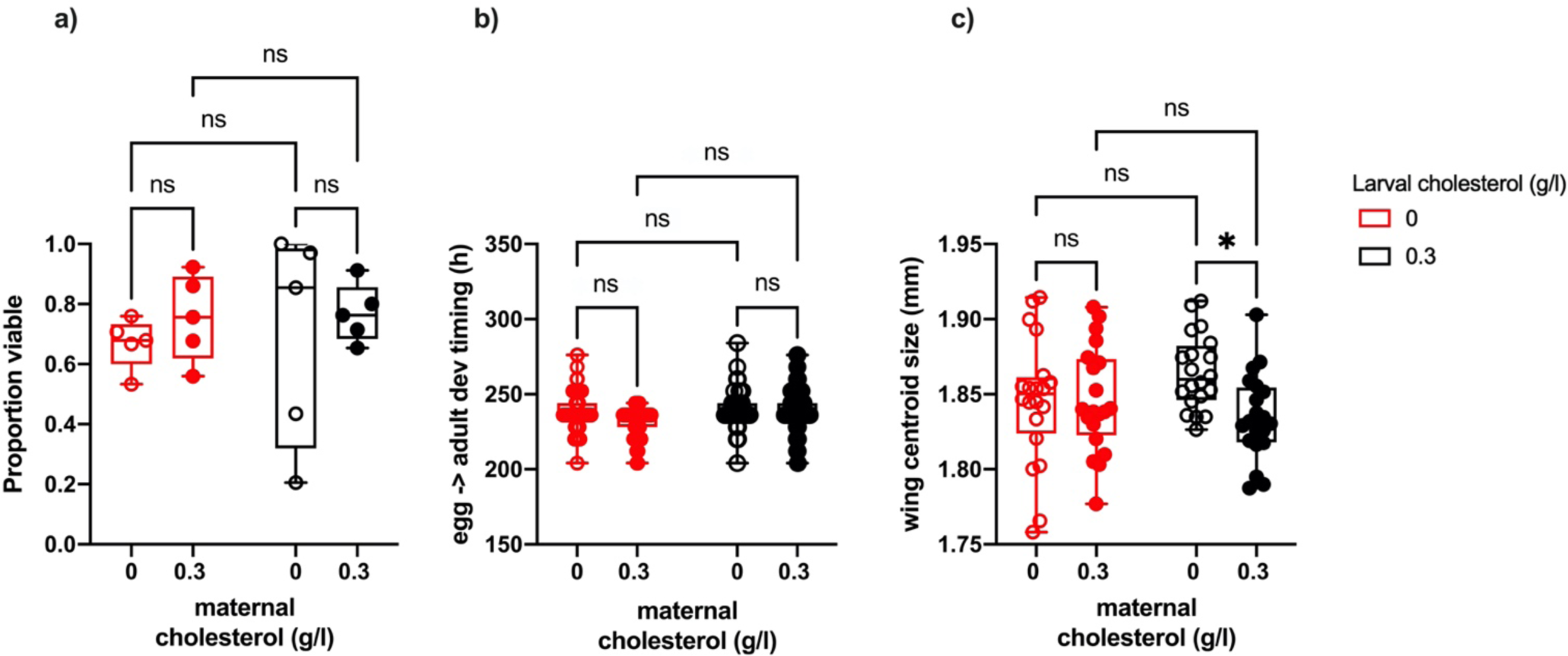
Egg-to-adult viability and egg-to-adult development timing were not significantly different across treatments (a-b). There was however a significant negative effect on offspring body size when both the maternal and larval diets were supplemented with cholesterol (in addition to naturally occurring sterols present in yeast) (c). Ns = non-significant, * = < 0.05, ** = < 0.005, *** = < 0.001

These findings indicate that the responses of the traits to cholesterol supplementation on the holidic diet (Figure 2) are not apparent when some dietary sterols are available to the mother (Figure 3) – even though those levels that are available are insufficient to support full lifespan (Zanco *et al*., 2021). Together, these data indicate that mothers prioritise sterols for use in producing viable eggs rather than maintaining the mother’s soma, supporting the Nutrient Recycling Hypothesis.

## Discussion

The mechanism by which DR increases animal lifespan is often attributed to the differential allocation of resources between reproduction and somatic maintenance (the disposable soma theory) (Kirkwood, 1977; Shanley and Kirkwood, 2000). Specifically, reproduction is repressed upon dietary restriction to redirect scarce nutrients to somatic maintenance, which increases the animal’s chances of surviving and reproducing at a later date. Here we show that during sterol limitation, the longer lifespan of flies during DR relative to those on higher nutrient diets cannot be explained by a strategic investment of the lifespan-limiting nutrient (sterols) into somatic maintenance and away from reproduction. Instead, our data indicate that flies constitutively invest sterols into reproduction to the maximum extent possible, which enhances early offspring viability. This is supported by the fact that any apparent enhancement of fitness as a result of sterol supplementation to the larval diet is only evident when mothers experience severe sterol limitation. Thus, flies on DR are longer lived because their commitment to reproduction is reduced, which spares the mothers from an early death caused by sterol depletion.

### Flies prioritise high quality egg production when fed a nutritionally deficient diet

In recent work to advance the Disposable Soma theory, the Nutrient Recycling Hypothesis (Adler and Bonduriansky, 2014) proposed that animals on restricted diets will attempt to utilise all available nutrients to maximise reproductive output rather than allocating it to somatic maintenance to prolong survival. The reason for employing this strategy is that prolonging survival is not likely to be advantageous when the chances of dying from extrinsic hazards are high, as is the case for most organisms in the wild. Our data support this concept since they indicate that female flies prioritise their supply of sterols to sustain high-quality reproduction whatever the cost to maternal lifespan. Flies likely maintain this level of optimal fecundity by first utilising as many resources from the diet as possible and secondly, by supplementing it from sterols stored in body tissues (Heier *et al*., 2021). Such a concept is curious as it suggests that DR may alter lifespan by stalling a program of tissue repurposing, which is in stark contrast to traditional models of ageing that envisage death as a side effect of random molecular damage (Kirkwood, 2005). Thus, we propose that the mechanisms for lifespan responses to DR in *Drosophila* are different from the mechanisms of ageing. This observation is consistent with the data of Mair *et al*. (2005) who showed that DR modifies a reversible risk of dying at any given age, and that this risk is different from, and additive to, the underlying, irreversible risk of dying that increases with advancing age (i.e. ageing). Importantly, our data also indicate that the mechanisms leading to earlier death on nutrient-rich diets should proceed in a reproducible (programmed) fashion; thus, if we can identify the process that causes lethal damage resulting from sterol depletion, we will discover the mechanism by which DR enhances *Drosophila* lifespan.

## Methods

### Fly Husbandry

All experiments were conducted using a wild type *Drosophila melanogaster* strain called Dahomey (Mair *et al*. 2005). These flies have been maintained in large numbers with overlapping generations to maintain genetic diversity. Flies were reared, mated prior to experiments and maintained under the same conditions described in Zanco *et al*. (2021).

### Experimental Diets

#### Holidic medium

To examine the effects of maternal cholesterol on developmental traits a fixed protein (amino acid): carbohydrate (sucrose) ratio of 1:3.6 was chosen (Zanco *et al*., 2021). This diet was made using the holidic medium described in Piper *et al*. (2014), in which free amino acids are used to make up protein equivalents. In this case, an amino acid ratio matched to the exome of adult flies (Flyaa) was utilised (Piper *et al*., 2017; Ma *et al*., 2020; Zanco *et al*., 2021). One of two cholesterol concentrations (0 and 0.3g/l) (Glentham Life Sciences, GEO100, #100IEZ) were then added to the otherwise identical media. We used cholesterol in the diet as opposed to ergosterol, because it is easily accessible, and has been shown to support *Drosophila* adult nutrition to the same extent as a yeast-based diet (Piper et al., 2014).

#### Yeast based diets

To examine the effects of both maternal and larval cholesterol on developmental traits using a standard laboratory medium, a fixed protein sugar/yeast (SY) diet was created using 50 g/l of sucrose (Bundaberg Sugar, Melbourne Distributors) and 100 g/l whole yeast autolysate (MP Biomedicals, LLC, #903312), previously described in Mair *et al*. (2005) and Zanco *et al*. (2021). We then added cholesterol (Glentham Life Sciences, GEO100, #100IEZ) at a concentration of either 0 or 0.3 g/l to both maternal and larval diets. This gave us a total of two maternal diets and two larval diets. Cholesterol was added as a powder which was mixed in with the sugar and yeast prior to cooking.

### Developmental trait assays

#### Egg-to-adult viability assay

Females were mated for two days post eclosion before being placed in vials (FS32, Pathtech) containing 3mL of treatment food with 5 replicate vials per diet and ten flies per vial. Diets utilised for this experiment included the holidic (fully defined synthetic) diet described above. Flies were transferred onto fresh food daily for 16 days so that egg and viability scores could be conducted. Eggs were imaged using a Zeiss Axiocam ERc 5s dissecting microscope and then counted manually. Eggs were then transferred to a standard sugar/yeast medium and left to develop (five vials of 10 eggs each). Egg-to-adult viability was scored as the number of eclosed adults.

#### Egg-to-adult viability, embryo to adult development timing and final body size assays

Females were mated for two days post eclosion and placed in vials (FS32, Pathtech) containing 3mL of treatment food with 10 replicate vials per diet and ten flies per vial. Diets utilised for this experiment included the holidic (fully defined synthetic) diets described above and the sugar/yeast diets described above (Bass *et al*. 2007; Piper *et al*. 2014; Katewa *et al*. 2016). Flies were transferred to fresh vials every two to three days. On day 10 (from the start of the experiment), adult flies were anaesthetised using CO_2_ and transferred to vials containing 3mL of our standard SY food (Bass *et al*. 2007; see below) either with or without cholesterol supplementation (0g/l vs. 0.3g/l) for five hours to lay eggs.

After laying, adults were discarded and eggs were imaged immediately using a Zeiss Axiocam ERc 5s dissecting microscope. Eggs were then counted manually. Scoring for developmental timing began on day 6 from egg lay and was done every 8 hours thereafter. Newly emerged flies were recorded and then stored in glycerol for wing analysis. To assess egg-to-adult viability, the number of eclosed adults were transferred out of experimental vials and counted. Viability was measured by allocating each individual a score of 1 if they survived to eclosion or 0 if they died. The assay continued until four consecutive days passed without any new emergences.

Wing size was calculated as a proxy for body size (David *et al*., 1997). The left wing of 20 individuals per treatment was dissected and mounted on a glass microscope slide (Clemson *et al*., 2016). Wing centroid size was calculated following the methodology described by (Clemson *et al*., 2016)

### Statistical analyses

All statistical analyses were performed using R (version 3.3.0, available from http://www.R-project.org/) and Graphpad Prism (version 8.4.2). Linear mixed effect models included egg count as a covariate, maternal and larval cholesterol levels as fixed effects and replicate vial as a random effect. The emmeans package was also used for pairwise comparisons, except for pairwise comparison of the daily viability data, in which case a two-way ANOVA was applied using Graphpad Prism (version 8.4.2). Plots were made in Graphpad Prism (version 8.4.2).

